# Concurrent neuroimaging and neurostimulation reveals a causal role for dlPFC in coding of task-relevant information

**DOI:** 10.1101/2020.04.22.054742

**Authors:** Jade B. Jackson, Eva Feredoes, Anina N. Rich, Michael Lindner, Alexandra Woolgar

**Affiliations:** MRC Cognition and Brain Sciences Unit, University of Cambridge, Cambridge, CB2 7EF, UK; Perception in Action Research Centre, Department of Cognitive Science, Macquarie University, Sydney, NSW 2109, Australia; School of Psychology and Clinical Language Sciences, University of Reading, Reading, RG6 6AH, UK

## Abstract

The way in which the brain prioritises processing of information relevant for our current goals is widely contested. Many studies implicate the dorsolateral prefrontal cortex (dlPFC), and propose that it drives brain-wide focus by biasing processing in favour of relevant information. An alternative, however, is that dlPFC is involved in the inhibition of irrelevant information. Here, we address this longstanding debate using the inferentially powerful approach of applying transcranial magnetic stimulation during functional magnetic resonance imaging (concurrent TMS-fMRI) and testing for changes in information coding using multivariate pattern analysis (MVPA). We ask whether dlPFC plays a causal role in prioritising information processing, and whether this is through *selection* of relevant information or *inhibition* of irrelevant information. Participants attended to one object feature whilst ignoring another feature of the same object. We reasoned that, if dlPFC is necessary for *selection*, active (disruptive) TMS should *decrease* coding of attended information compared to the low intensity (control) condition. Conversely, if right dlPFC is crucial for *inhibition*, active TMS should *increase* coding of irrelevant information relative to the control condition. The results showed that active TMS decreased coding of *relevant* information throughout the frontoparietal multiple demand regions, and that this impact was significantly stronger than the effect of TMS on *irrelevant* information coding, which was not statistically detectable. These data provide causal evidence for a specific role of dlPFC in supporting the representation of task-relevant information and demonstrate the crucial insights into high level cognitive-neural mechanisms possible with the combination of TMS-fMRI and MVPA.

## Introduction

A critical aspect of successful goal-directed behaviour is the ability to distinguish between information that is relevant for your current task and distracting irrelevant information. How does the brain achieve this selection? One prominent theory (“adaptive coding hypothesis”; Duncan, 2001) posits that prefrontal neurons adjust their responses to code the information relevant for behaviour, biasing and modulating responses in specialised cortices (e.g. Desimone & Duncan, 1995; Corbetta & Shulman, 2002). In a similar vein, Miller and Cohen (2001) suggest that attention biases competing inputs in favour of relevant information (see also Desimone, 1998). Evidence for these models stem from non-human primate research where prefrontal neurons are shown to maintain task-relevant information in delayed-response tasks (e.g. Fuster & Alexander, 1971; Fuster, 1973) and flexibly encode the behavioural significance of visual stimuli, regardless of their physical properties (Freedman, Riesenhuber, Poggio, & Miller, 2001; Cromer, Roy, & Miller, 2010; Roy, Riesenhuber, Poggio, & Miller, 2010; Stokes et al., 2013).

In the human brain, a network referred to as the multiple-demand (MD) network encodes a range of task features (Woolgar, Jackson, & Duncan, 2016) with a strong preference for attended information over information that is irrelevant (Erez & Duncan, 2015; Woolgar, Williams, & Rich, 2015; Jackson, Rich, Williams, & Woolgar, 2017; Jackson & Woolgar, 2018). This network includes the dorsolateral prefrontal cortex (dlPFC), the anterior insula and frontal operculum (AI/FO), intraparietal sulcus (IPS), and the pre-supplementary motor area and adjacent anterior cingulate (ACC/pre-SMA). Of these regions, the dlPFC is frequently emphasised as the likely candidate for top-down signals that influence neural activity according to behavioural relevance (e.g. Knight, Staines, Swick, & Chao, 1999; Shimamura, 2000; Curtis & D’Esposito, 2003; Miller & D’Esposito, 2005). This view stems from the non-human primate literature, as well as human functional magnetic resonance imaging (fMRI) data. For example, blood flow increases in this region during tasks requiring selection of relevant over competing irrelevant information compared to neutral trials (e.g. Milham, Banich, & Barad, 2003). Thus, there is evidence suggesting that the MD network, and the dlPFC in particular, plays a role in attentional selection and resultant top-down modulation.

Two mechanistic possibilities for how top-down influences could be achieved are via the enhancement of task-relevant information, and/or by the inhibition of irrelevant information (Kanwisher & Wojciulik, 2000; Aron, 2007). Despite decades of research, the current literature is at an impasse. One reason for this is because neuroimaging work is unable to draw *causal* links between network interactions, brain function, and behaviour. Here, we addressed this debate using a combination of concurrent transcranial magnetic stimulation (TMS) and fMRI with multivariate pattern analyses (MVPA). The disruptive effect of TMS allows a test of causality rather than only correlation, and the use of MVPA allows inference about the information coded in the system, rather than simply whether blood flow increases or decreases (Lewis-Peacock & Postle, 2012). Thus, together, our approach can test for the causal influences of dlPFC on information representation across the brain.

During fMRI acquisition, participants attended to one feature (colour or form) of an object whilst ignoring an irrelevant feature of that same object (Figure 1). On each trial we applied a short train of either high (Active) or low (Control) intensity TMS stimulation to right dlPFC (Figure 2). We asked whether the dlPFC (1) upregulates relevant information (e.g. Desimone, 1998); (2) downregulates irrelevant information (e.g. Shimamura, 2000); or (3) a combination of both. Right dlPFC was selected as the stimulation target over left dlPFC as TMS to this region of interest (ROI) has been shown to modulate activation in a task requiring selective attention (Feredoes, Heinen, Weiskopf, Ruff, & Driver, 2011). If right dlPFC normally *enhances* coding of attended information, then disrupting it with TMS should *decrease* coding of attended stimulus features across the MD network (disruption to the system) and visual cortices (top-down modulation). Conversely, if the right dlPFC *suppresses irrelevant* information, then disrupting this function should *increase* coding of the *unattended* feature. If right dlPFC plays both roles, then Active TMS should both *decrease relevant* and *increase irrelevant* information coding. Another possibility is that the right dlPFC may play a general role in supporting all information processing (no specific role in attentional selection). Under this alternative, Active TMS would decrease both relevant and irrelevant information coding. A final possibility is that the right dlPFC has no role in supporting information processing, in which case we would expect no change in information coding under Active versus Control TMS. We tested these predictions in the MD regions, to test network function (Duncan, 2010), and in visual cortices (lateral occipital complex (LOC), V4, early visual cortex) and across the whole brain (using a roaming searchlight) to assess top-down modulation (e.g. Desimone & Duncan, 1995).

**Figure 1:**
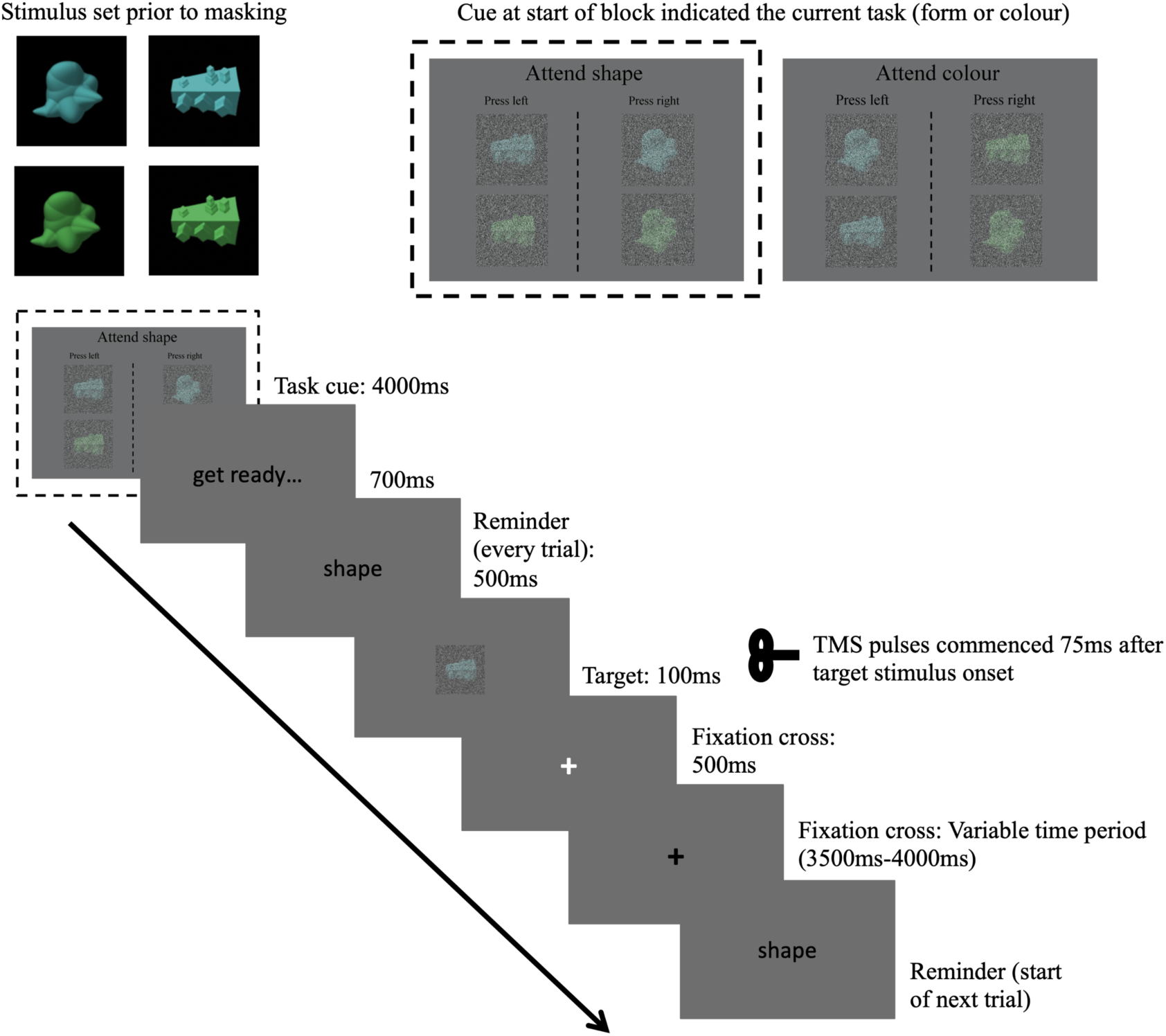
Main concurrent TMS-fMRI task. On each trial, participants saw a target object which they had to categorise according to either its form (cuby/smoothy) or colour (green/blue) in alternating blocks. A picture cue at the start of each block indicated the current task context (form or colour; upper right inset). On each trial a cue reminded participants of the current task (500ms) followed by the object to categorise (100ms), during which they received a train of 3 TMS pulses (13Hz; high or low intensity, onset 75ms after stimulus). Participants were instructed to respond before the white cross (500ms) turned black. If participants responded within the 500ms, the white cross turned black for the remainder of the 500ms and was in either case followed by a black cross for a jittered interval of 3500ms-4000ms. In the example shown, participants are cued to attend to shape and the correct response would be the left button. The stimulus-response mappings for the two dimensions meant that for some stimuli its shape and colour required the same button response (congruent) and for other stimuli its shape called for one button and its colour called for the other button (incongruent). The trial depicted in this figure is a congruent trial as the correct button response for its shape is left, and its blue colour also indicate the left button during the colour task. On the upper left of the figure, the four presented objects are displayed prior to masking for illustration purposes.

**Figure 2:**
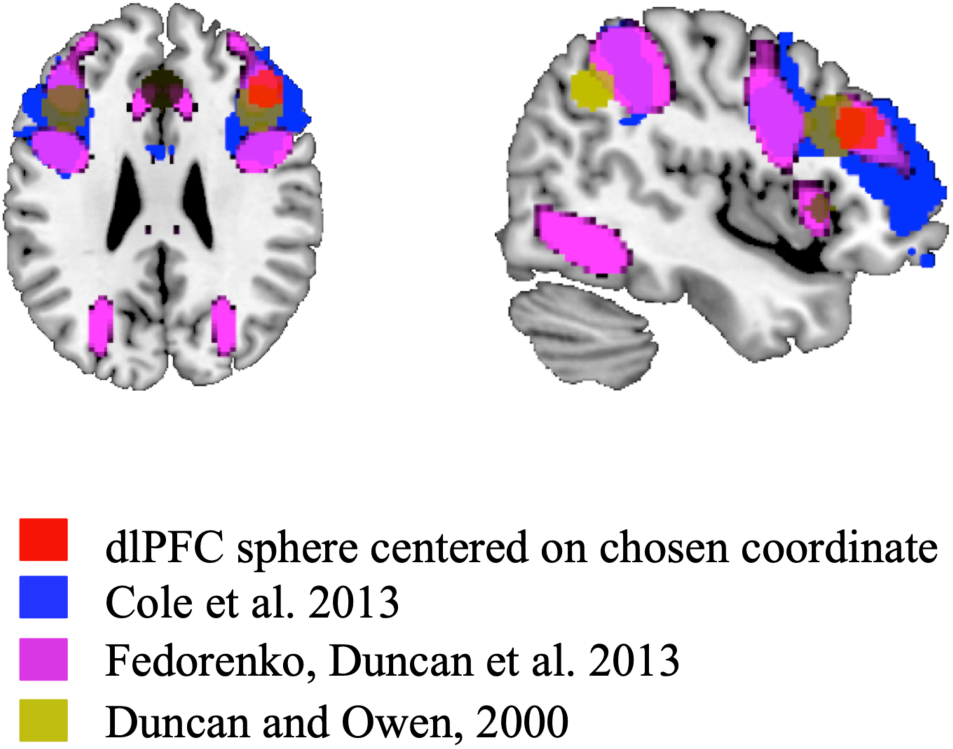
Selection of TMS target site (right dlPFC). Overlaid functional activation and connectivity maps from previous studies (as in legend) were used to select MNI152 coordinates [44 31 28] as a group level reference for dlPFC (visualised here as a red sphere of radius 8mm). After deforming to native space, we defined the target stimulation site on an individual-participant basis using the peak activation from individual-participant functional localiser data that was within 14mm of this reference point.

## Methods

### Participants

Thirty-one healthy volunteers signed up for the experiment. However, four participants did not pass the TMS screening requirements, and seven participants did not complete the second scanning session, so their data are not included. The final group consisted of twenty participants (15 females, 5 males; mean age = 21.6 years, SD = 3.36). All participants were right-handed with normal or corrected-to-normal vision and no history of neurological or psychiatric disorder. Participants gave written informed consent and received £30.00. The experiment was approved by the University of Reading Research Ethics Committee.

### Stimuli

Stimuli were abstract novel “smoothy” and “cuby” objects created using custom scripts (Op de Beeck, Baker, DiCarlo, & Kanwisher, 2006). The stimulus set consisted of 4 objects (see Figure 1) which were either blue (RGB: 98 179 180) or green (RGB: 95 171 96) and were one of two novel shapes (“cuby” or “smoothy”). To increase perceptual difficulty, we superimposed a noise filter in Adobe Photoshop CC (2014), which applies random pixels to the picture. The colour values used for the noise creation were distributed on a bell-shaped (i.e. Gaussian) curve. We applied the monochromatic filter which allows for a degree of colour preservation within the image, because the filter applies pixels that differ only in tone, and not in actual colour, from the original image. In separate blocks of trials, participants reported either object colour (blue or green), or object form (cuby or smoothy). Thus, the visual feature that was relevant varied between blocks. We controlled stimulus presentation with a PC running the Psychophysics Toolbox-3 package (Brainard, 1997) in MATLAB (Mathworks).

### Procedure

Participants attended two sessions separated by 2-8 days. In Session 1, we determined the participant’s resting motor threshold (MT) and familiarised them with the sensation of TMS. They also had a structural MRI scan and completed three functional localiser tasks in the MR scanner to determine the stimulation site and ROIs for further analysis. In Session 2, participants completed the main task in the scanner, with concurrent TMS. The same TMS machine and coil were used in Session 1 for determining individual MTs as in Session 2 for the main experiment. Below we outline the procedures in detail.

#### Session 1

For each participant, we first acquired their resting MT outside of the scanner. For this we determined the minimum intensity at which a single pulse through the TMS coil, positioned over the hand area of the primary motor cortex, produced a visible twitch in the abductor pollicis brevis when at rest, in 5 of 10 successive pulses. Individuals’ MTs determined stimulation intensity for that participant in Session 2. Stimulation intensity in the scanner in Session 2 was pseudo-randomly varied over trials within a block at either 110% (Active stimulation) or 40% (Control stimulation) of the individual participant’s MT. After their MT was determined, participants were given instructions and completed the structural scan and localiser tasks.

##### Localiser: Right dlPFC

We used a functional localiser alongside activation maps from previous studies (see section: Selection of the stimulation target) to determine our target site in dlPFC. The functional localiser was a modified version of the main experimental task. On each trial, participants saw an object which they had to categorise according to either its form (cuby/smoothy) or colour (green/blue) in alternating blocks. Within each block there was a mini block of 8 congruent followed by 8 incongruent trials (or *vice versa*, for the localiser, congruent and incongruent trials were blocked to give maximal power for analysis). Congruent trials comprised objects where the response button for the irrelevant dimension was congruent with the required response for the relevant dimension. Conversely, the incongruent blocks consisted of objects where the response for the irrelevant dimension was incongruent with the required response for the relevant dimension. For example, if a participant was instructed to respond ‘left’ for blue and ‘right’ for green when reporting colour, and ‘left’ for cuby and ‘right’ for smoothy when reporting form, then a green cuby would be an incongruent trial (i.e. ‘right’ button response in colour task, ‘left’ button response in form task) and a blue cuby would be congruent. Rest blocks were also included (black cross at fixation: 16s) in between the colour and form alternating task blocks. Task order (colour/form) and block type (congruent/incongruent) were counterbalanced across participants.

Participants practiced outside the scanner for a minimum of 6 blocks and until they achieved > 70% performance. For the first 2 practice blocks and after each stimulus presentation, participants viewed a black cross (2000ms) during which time they were told that their responses were recorded. Following the first 2 blocks and to mimic the main experimental task, immediately following object presentation, a white cross appeared for 500ms followed by a black cross for 1100ms. If participants responded within the 500ms, the white cross turned black for the remainder of the 500ms. The white cross served to add time pressure, as participants were told that their responses were only recorded during the white cross period (responses were, however, still recorded during the black cross period). In the first two practice blocks participants received feedback on every trial. Following this, they only received feedback (percent correct) at the end of the block.

In the scanner, participants completed 2 localiser runs (6 minutes each). Within each run there were 18 blocks of each condition: congruent (16.8s, 8 trials/block), incongruent (16.8s, 8 trials/block), and rest (16.8s). Participants received feedback (percent correct) at the end of every block. In the second run, the button response mapping (the left- and right-hand button response associated with each colour/object) was switched to mimic the procedure of the main task (below). The response mapping switch was included to dissociate activity associated with participant’s motor responses from that with the stimulus features. The button response mapping order was counterbalanced across participants.

We defined dlPFC as the region close to the inferior frontal sulcus that responded more strongly on incongruent than congruent trials (see “Selection of the stimulation target” below). We reasoned that this activity would reflect the processes required to selectively attend to relevant over irrelevant information.

##### Localiser: LOC

The LOC localiser was designed to functionally identify object-sensitive cortex. Participants viewed centrally located intact and scrambled versions of black and white objects in 16.8s blocks of 16 trials (1100ms / trial), whilst attending to a central fixation cross. Participants indicated via a button response when the fixation cross changed from black to blue. There were 21 blocks consisting of alternating blocks of whole objects, scrambled objects, and rest blocks (order counterbalanced across participants). The EPI time was 6.25min.

##### Localiser: Early visual cortex

The final localiser was designed to identify the region stimulated by visual information at fixation (encompassing the same area of central visual field as the objects in the main experimental task). Participants viewed a small white circle at the centre of the screen that was visible across all blocks. Rest blocks (16s) consisted only of the circle presentation. The stimulus blocks (16s) included either a flickering checkerboard presented only at fixation (the spatial extent matched the objects in the main task) or a checkerboard filling the entire field of view leaving only a blank grey screen in the place of the fixation checkerboard. Participants were required to press a button when the central circle on the screen flashed green (0.1s) to encourage attention to the screen. The EPI time was 8 mins.

#### Session 2

In Session 2, we implemented the Brainsight infrared frameless stereotaxy neuronavigational system (Rogue Research Inc.) to navigate to each participant’s individual target stimulation location, and marked the site on their scalp. Following this, participants practiced the main task outside the scanner and then completed 8 runs in the scanner with simultaneous online TMS on every trial.

##### Selection of the stimulation target

We defined the target stimulation site based on individual subject data cross-referenced to functional activation (those regions showing activation for a wide range of tasks from Duncan & Owen, 2000; Fedorenko, Duncan, & Kanwisher, 2013)) and connectivity data (the frontoparietal network from Cole et al., 2013)) from the literature (Figure 2).

We calculated two contrasts from our dlPFC localiser: (1) *incongruent > congruent;* and (2) *(incongruent + congruent) > rest*. We then derived a “comparison sphere” centered on MNI co-ordinates [44 31 28] at the intersection of the activation and connectivity maps and deformed it into individual subject native space. If the dlPFC localiser peak activation (see section “Localiser: Right dlPFC” for contrast details) from contrast (1) was less than 14mm from the centre of the comparison sphere, then the central coordinate of peak activation was selected for that participant as their stimulation target (n=12; furthest individual peak coordinate was 13.08mm). If contrast (1) showed no activation (minimum initial threshold of p < 0.001, uncorrected) within range of the comparison sphere, then we compared activation for contrast (2) applying the same procedure (n=5). For the remaining 3 participants, there were no clusters of activation from either contrast within 14mm of the comparison sphere, so the central point of the sphere was chosen as the target.

##### Neuronavigation

We used the Brainsight neuronavigational system to guide coil placement to the individual stimulation target coordinates, using standard neuronavigation routines. The target location was marked on the scalp for stimulation inside the scanner. The final position of the TMS coil was adjusted in the scanner to fit within the MR head coil and to be comfortable for participants. The TMS coil was oriented with the handle pointing posteriorly with respect to the participant’s head, and roughly parallel to the midline, to target the frontoparietal network as opposed to the default mode network (Opitz, Fox, Craddock, Colcombe, & Milham, 2016). For some participants, adjustments to the coil orientation were necessary to ensure no part of the TMS coil was touching the head coil. Orientation adjustments were also made, if necessary, to minimise any discomfort produced by the stimulation, which can sometimes be the case for the dlPFC scalp location.

##### Concurrent TMS-fMRI (main experimental task)

Before entering the scanner, participants again practiced the task for at least 6 blocks without TMS. For this and the main task, the task was blocked (colour, form). Within each block congruent and incongruent trials were presented *pseudo*-randomly (not in mini-blocks as they had been for the localiser). There were no rest blocks and the timings were slowed down, relative to the task localiser (see Figure 1 for task design). This was to ensure that the time between the TMS trains met with safety recommendations (Rossi, Hallett, Rossini, Pascual-Leone, & Group, 2009). During the practice, participants received feedback after each trial for the first two blocks as well as feedback (percent correct) at the end of the block. For the last four blocks of practice, and in the scanner, participants only received feedback at the end of the block. Participants repeated the last four blocks of practice trials until they scored > 70% correct.

In the scanner, participants completed 8 runs (6.3 minutes each) of the task with concurrent TMS. For this, an MR-compatible TMS figure-8 stimulating coil (MRI B90 II, MagVenture, Farum, Denmark) was held firmly in position inside the MR head coil, by a custom-made non-ferromagnetic coil holder. The cable of the TMS coil passed through the back of the scanner and out through a wave guide and connected to the TMS machine (Magpro X100, MagVenture, Farum, Denmark) located in the MR control room. In each run participants completed one block of the colour task and one block of the form task. The block started with a picture cue (4000ms) indicating the current task context and response mapping. On each trial participants first saw a written cue reminding them of the current task (“colour” or “shape”, 500ms) followed by the target object (100ms). Participants received a train of three TMS pulses: the first pulse at 75ms after target stimulus onset, and then the remaining pulses separated by 75ms (13Hz). A TMS protocol of 3 pulses at 13Hz was chosen here as inhibitory effects on behaviour have been observed in previous work with a similar protocol (e.g. Postle et al., 2006; Feredoes, Tononi, & Postle, 2007). Pulses were triggered by a MATLAB (Mathworks) script on a PC which also received pulse timings from the MR machine and controlled the visual stimulus delivery to the projector. Pulses were delivered to coincide with the readout phase of slice acquisition. They were timed so that the artifact caused by each TMS pulse would affect only one MRI slice and would occur at a different slice of each volume (Bestmann, Baudewig, & Frahm, 2003). The affected slice was later discarded and interpolated over (see below). The train of pulses was delivered at 110% (Active) or 40% (Control) of participant’s MT, varying pseudo-randomly over trials. There were a total of 1536 pulses across all of the runs, complying with published safety limits for TMS stimulation (Rossi, Hallett, Rossini, & Pascual-Leone, 2009). Immediately following object presentation (100ms), a white cross appeared for 500ms followed by a black cross for a variable time-period (3500-4000ms). If participants responded within the 500ms, the white cross turned black for the remainder of the 500ms. The participants were told that their responses were only recorded during the white cross period (responses were actually still recorded during the black cross period). Participants received feedback at the end of each block. After the first four runs, the button-response mapping for both colour and form was swapped (e.g., if the button response for green had been the left-hand button response, green would now require a right-hand button response). Mappings were swapped for both colour and form, so the congruency of response for any given object did not change.

**Figure 3:**
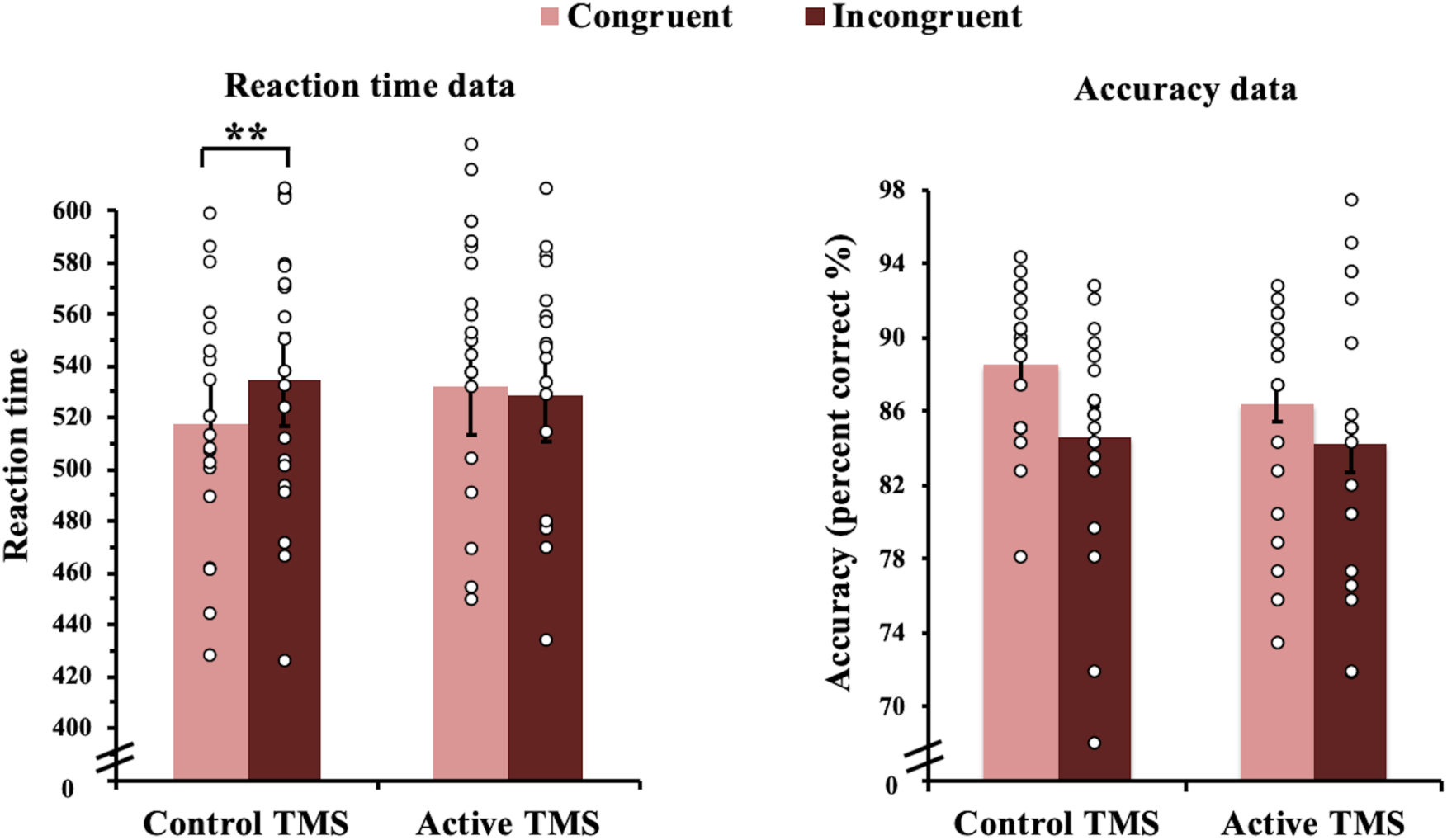
Reaction time (left panel) and accuracy data (right panel). RT data (correct trials only) revealed that participants were faster in congruent trials than incongruent trials under the Control TMS condition but showed no evidence of a congruency effect under Active TMS (significant interaction). Accuracy data showed a main effect of congruency (no interaction). Open circles represent individual subject data. Error bars indicate standard error. **p<0.01. N=20.

**Figure 4:**
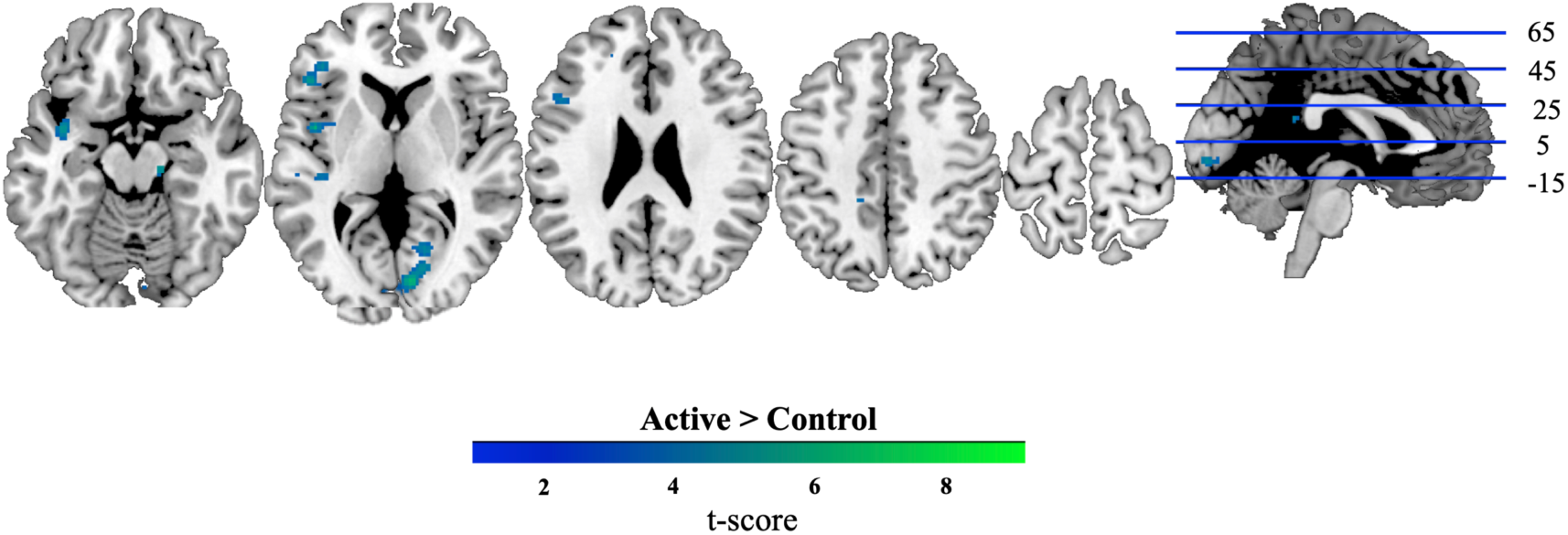
Brain regions showing a larger BOLD response under Active than Control TMS (univariate contrast). We examined differences in overall BOLD response under Active (high intensity) and Control (low intensity) TMS using a mass-univariate whole brain approach. Active and Control trials were modelled separately, and BOLD responses contrasted at the second level with paired t-tests at each voxel (for both Active > Control and Control > Active). Group-level analysis for Control > Active showed no significant clusters of activation at the set threshold. These results were thresholded at *p <* 0.0001 (FWE correction *p* < 0.05 at cluster level). Coordinates of peaks are given in Table 1. N=20.

**Figure 5:**
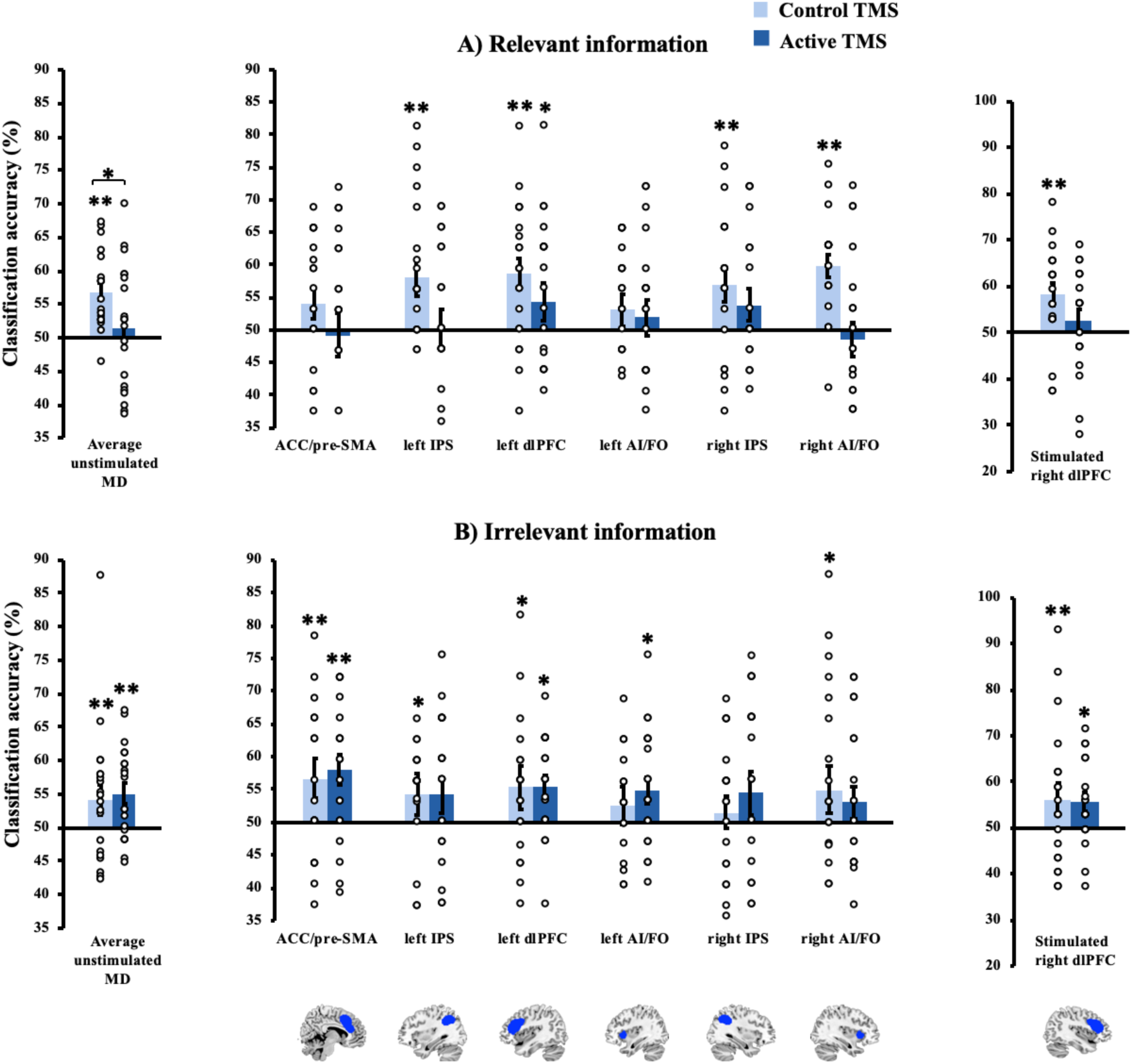
Decoding results in MD regions (left: average across unstimulated MD regions; centre: unstimulated MD regions separately; right: stimulated right dlPFC). Upper panel shows coding of relevant information under Control and Active conditions, collapsed across feature (colour, form). Lower panel shows coding of irrelevant information under Control and Active conditions, collapsed across feature. All bars represent coding of identical stimulus information, variation in the strength of coding is driven by TMS intensity and whether the information was relevant for the participant’s current task. Open circles represent individual subject data. The ANOVA on the unstimulated MD regions (left panels) showed stronger coding of relevant features across MD regions under Control compared to Active TMS, but there was no modulated coding by TMS on irrelevant information (BF_10_ = 0.08). Due to outliers (>3 SD from the condition mean) we performed a log transformation on the unstimulated MD region data before calculating ANOVAs and permutation tests against chance. The data displayed are in the untransformed form prior to log transformation. The ANOVA for right dlPFC (factors: *TMS, Feature*, and *Relevancy*; Right panels) showed no main effects or interactions. Error bars indicate standard error. The significance markings for individual bars indicate whether coding was significantly greater than chance in each condition separately (by permutation). *p<0.05, **p<0.008 (corrected for multiple comparisons). N=20.

**Figure 6:**
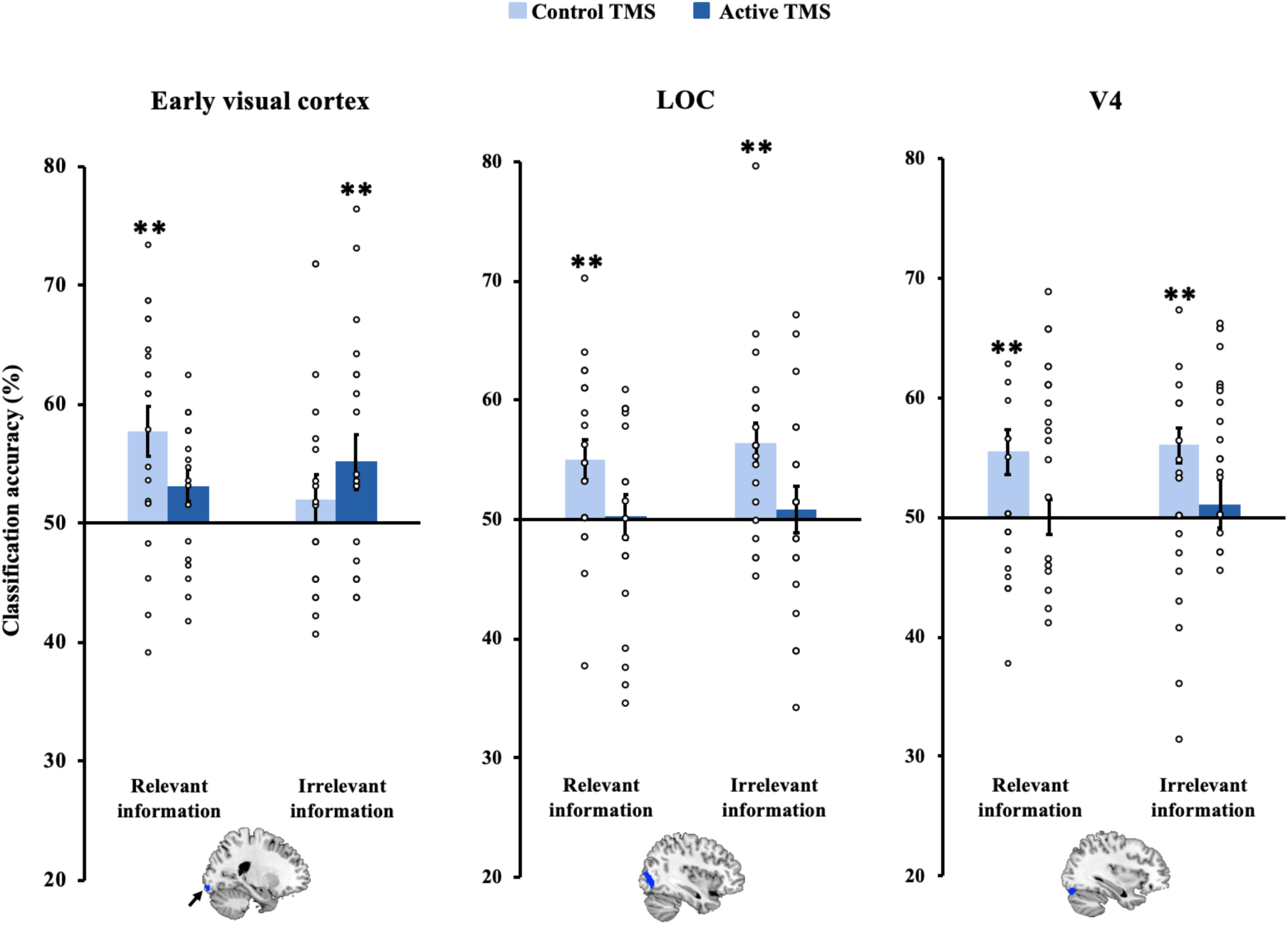
Coding in visual ROIs under Control and Active TMS. Left panel. Early visual cortex (central visual field). This ROI was derived from individual-participant localiser data and defined as the region stimulated by visual information at fixation (encompassing the same area of central visual field as the objects in the main experimental task) minus visual information outside fixation. There were no significant main effects or interactions. **Centre panel:** Lateral Occipital Complex (LOC). This ROI was derived from localiser data as the region more active for viewing of whole objects over scrambled objects. There was stronger coding under the Control TMS condition compared to the Active condition modulated by a Feature*TMS interaction reflecting a stronger effect of TMS on colour than form coding. **Right panel:** V4. This ROI was derived from coordinates from the literature (Van Leeuwen, Petersson, Langner, Rijpkema, & Hagoort, 2014) and transformed into native space for each participant. There was again a main effect of TMS modulated by a Feature*TMS interaction. Error bars indicate standard error. Significance markings for individual bars indicate whether coding was significantly greater than chance in each condition separately (by permutation). *p<0.05; **p<0.01. Individual data points are marked on bars. N=20.

**Figure 7:**
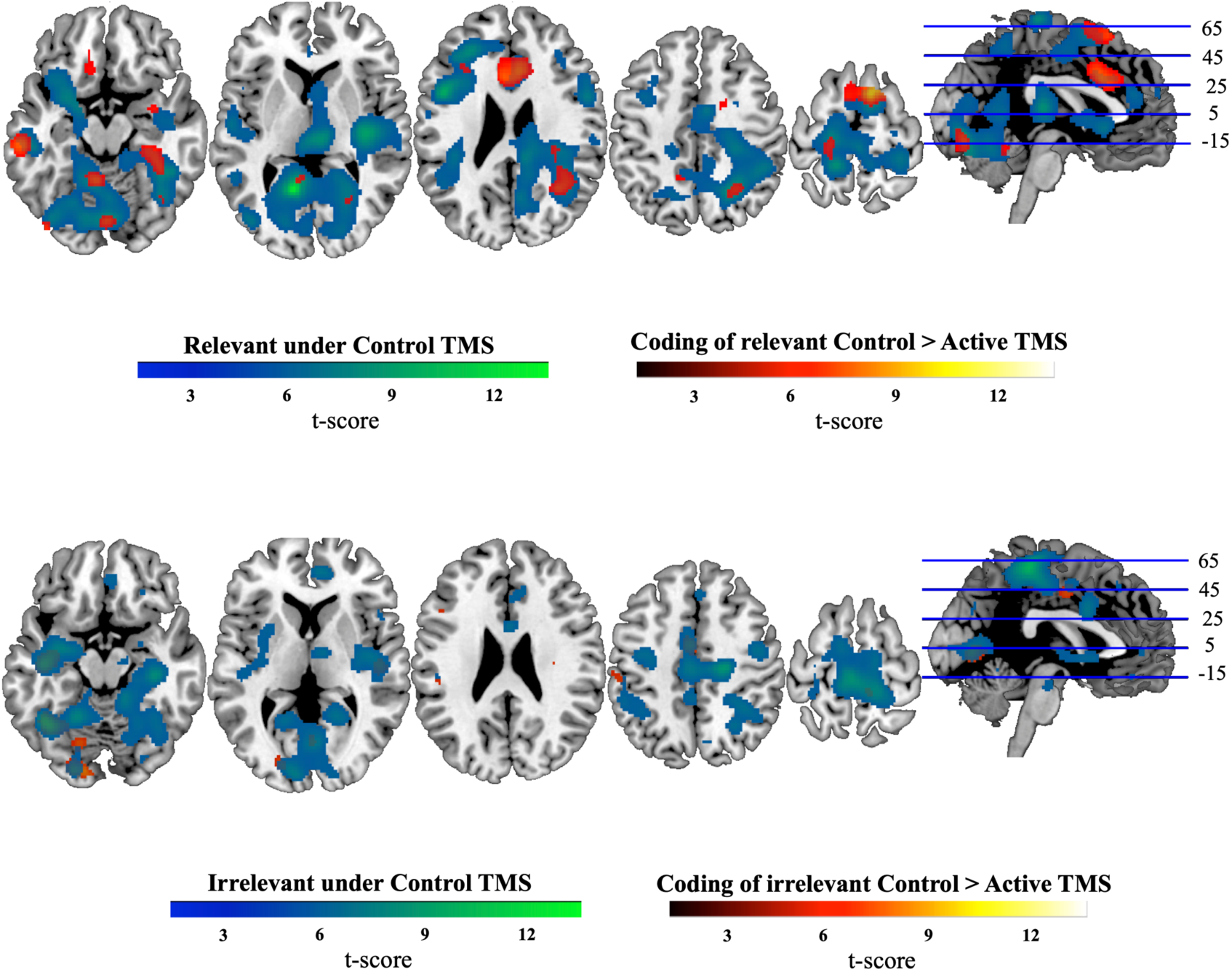
Information coding assessed with a roaming searchlight (relevancy maps). Whole brain maps where patterns of activation in the local neighbourhood (10 mm sphere) discriminated relevant information under Control TMS (upper panel, blue-green), relevant coding was significantly reduced from Control to Active TMS (upper panel, red-yellow), patterns discriminated irrelevant information under Control TMS (lower panel, blue-green), and where irrelevant coding was significantly reduced from Control to Active TMS (lower panel, red-yellow). No other contrasts showed significant clusters and are not depicted. Results were thresholded at *p <* 0.0001 (FWE correction at cluster level, *p* < 0.05). Coordinates of peak decoding are given in Tables 2, and 3. N=20.

**Figure 8:**
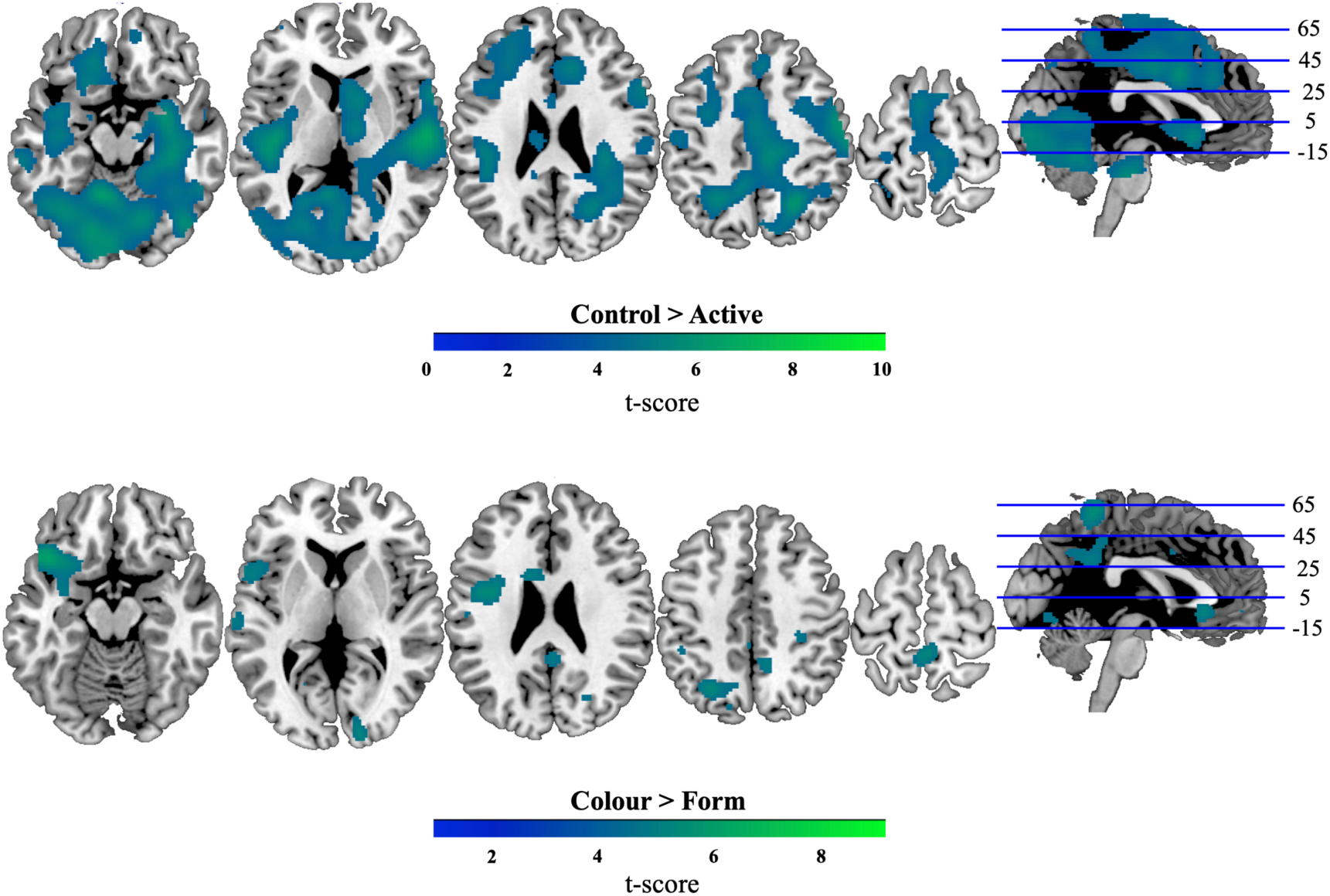
Information coding assessed with a roaming searchlight (repeated measures ANOVA). Whole brain maps where patterns of activation in the local neighbourhood (10 mm sphere) discriminated information more strongly under Control > Active (upper panel), and Colour > Form conditions (lower panel), as revealed by a three-way repeated measures ANOVA. Results were thresholded at *p <* 0.0001 (FWE correction of *p* < 0.05 at cluster level). N=20.

### Data acquisition

FMRI data were collected for both scanning sessions using a Siemens Magnetom Trio 3T whole-body MRI scanner at Centre for Integrative Neuroscience and Neurodynamics, Reading University, Reading, UK.

#### Session 1: Localisers

We used a sequential ascending T2*-weighted EPI acquisition sequence with the following parameters: acquisition time 2080ms; echo time 30ms; 60 oblique axial slices with a slice thickness of 3.0 mm and a 0.70 mm inter-slice gap; in plane resolution 3.0×3.0 mm; matrix 64×64; field of view 256 mm; flip angle 78°. T1-weighted MPRAGE structural images were also acquired for all participants (slice thickness 1.0 mm, resolution 1.0×1.0 mm).

#### Session 2: TMS-fMRI experiment

We used a sequential ascending T2*-weighted EPI acquisition sequence with the following parameters: acquisition time 2450ms; echo time 30 ms; 35 oblique axial slices with a slice thickness of 3.0 mm and a 0.70 mm inter-slice gap; in plane resolution 3.0×3.0 mm; matrix 64×64; field of view 256 mm; flip angle 90°; 50% phase oversampling in the phase encoding direction to shift any Nyquist ghost artefact, due to the presence of the TMS coil, to outside the volume of interest.

### Preprocessing

#### Session 1: Localisers

MRI data were preprocessed using SPM 5 (Wellcome Department of Imaging Neuroscience, www.fil.ion.ucl.ac.uk/spm) in MatLab 2013b. Functional MRI data were converted from DICOM to NIFTI format, spatially realigned to the first functional scan and slice timing corrected, and structural images were co-registered to the mean EPI. EPIs were smoothed (8 mm FWHM Gaussian kernel) and in all cases the data were high pass filtered (128s). Structural scans were additionally normalised to the T1 template of SPM5 (Wellcome Department of Imaging Neuroscience, London, UK; www.fil.ion.ucl.ac.uk), using SPM5’s segment and normalise routine. This was done to derive the individual participant normalisation parameters needed for transformation of ROIs into native space and TMS target definition, and to normalise the searchlight classification maps derived in native space.

#### Session 2: TMS-fMRI experiment

Following conversion of the functional data from DICOM to NIFTI format, we removed the slices that were affected by TMS pulses (one slice per pulse). We first identified slices with a signal magnitude of > 1.5 SD from the run mean and visually inspected them for presence of the TMS artefact. These slices were replaced by temporal interpolation of the signal values of the same slice from the preceding and succeeding volumes (following Feredoes et al., 2011). Next, we manually removed and interpolated over any remaining slices that were acquired during TMS pulse delivery, identifying them based on timing and visual inspection. This was necessary because, depending on the affected slice, the Control TMS condition did not always produce deviations > 1.5 SD from the mean. We ensured that the same number of slices were removed and interpolated over in the Active and Control TMS conditions. Aside from this slice removal step for Session 2 data, preprocessing followed the same steps as Session 1. EPIs from Session 2 were smoothed slightly (4 mm FWHM Gaussian kernel) to improve signal-to-noise ratio for multivariate analyses, and were smoothed separately with a larger smoothing kernel for univariate analyses (8 mm FWHM Gaussian kernel).

### ROI definition and statistical analysis

#### Univariate contrasts

##### Localiser: Right dlPFC

We used the multiple regression approach of SPM5 to estimate values corresponding to the congruent, incongruent and rest conditions. Blocks were modelled using a box car function lasting 16.8s convolved with the hemodynamic response of SPM5. The run mean was included in the model as a covariate of no interest. Whole-brain analyses (paired t-tests) compared blood-oxygen-level-dependent (BOLD) responses across the following conditions: [incongruent – congruent], which was the congruency effect, and [(incongruent + congruent) – rest], which was the task effect. The resulting map was thresholded such that there was at least one cluster with a minimum size of 20 voxels. We used this lenient thresholding as the purpose was to identify an ROI for target stimulation. The peak of one of these clusters was chosen as the target stimulation site as explained in the section: “Selection of the stimulation target”.

##### Localiser: LOC

We estimated values pertaining to the whole and scrambled object conditions. Blocks were modelled using a box car function lasting 16s convolved with the hemodynamic response of SPM5. The run mean was included in the model as a covariate of no interest. Mass univariate paired t-tests compared voxelwise BOLD response in the two conditions (whole objects minus scrambled objects). For each subject separately, the resulting maps were thresholded such that there was at least one cluster with a minimum size of 20 voxels in LOC in each hemisphere (close to coordinates ([-44 -67 -10], [41 -67 -11], and [-41 -78 -2], [39 -73 -3]), from previous studies; Grill-Spector et al., 1999; Grill-Spector, Kushnir, Hendler, & Malach, 2000). We used this lenient thresholding as the purpose was to identify an ROI for MVPA analysis. The mean size of LOC across participants was 150.8 voxels (S.D. 85.8).

##### Localiser: Early visual cortex

Blocks were modelled using a box car function lasting 16s convolved with the hemodynamic response of SPM5. The run mean was included in the model as a covariate of no interest. We contrasted BOLD responses with paired t-tests in the two conditions (fixation minus outside-fixation checkerboard). The resulting maps were thresholded so that there were two clusters, of minimum size of 20 voxels, in early visual cortex. The mean size of early visual cortex across participants was 84.3 voxels (S.D. 9.13).

##### The effect of TMS on overall activation levels

We examined differences in overall BOLD response under Control (low intensity) and Active (high intensity) TMS using a mass-univariate whole brain approach. A General Linear Model (GLM) was estimated for each participant using the realigned, slice-time corrected and smoothed normalised EPI images (8 mm FWHM Gaussian kernel) from Session 2. We modelled Control and Active trials separately and contrasted BOLD responses at the second (random effects across subjects) level with one-tailed paired t-tests at each voxel (for both Control > Active and Active > Control). The results were thresholded at *p <* 0.0001 (cluster-level family wise error (FWE) correction for multiple comparisons). All coordinates are reported in MNI152 space.

#### V4 (colour-responsive cortex) definition

The V4 ROI was defined from previous data (Van Leeuwen, Petersson, Langner, Rijpkema, & Hagoort, 2014), centred on coordinates [-32 -82 -20; left hemisphere] and [32 - 82 -20; right hemisphere]. The mean size of V4 across participants was 56.1 voxels (S.D. 5.23).

#### MD network definition

MD ROIs were defined using co-ordinates from a previous review of activity associated with a diverse set of cognitive demands (Duncan & Owen, 2000) using the kernel method described in Cusack, Mitchell, and Duncan (2010) as in our previous work (Woolgar, Hampshire, Thompson, & Duncan, 2011; Woolgar, Thompson, Bor, & Duncan, 2011; Woolgar et al., 2015; Jackson et al., 2017; Jackson & Woolgar, 2018). We opted to use a template definition of the MD network because recent work has indicated that there is no particular benefit to using individual subject functional localisers to define these regions for multivariate analysis (Shashidhara, Spronkers, & Erez, 2019), whereas the template approach made it easy to compile the data across the group. The average size in voxels across participants of ACC-pre SMA was 792.0 (S.D. 65.8), IPS; 787.8 (S.D. 26.1), left dlPFC; 676.8 (S.D. 43.8), right dlPFC; 685.6 (S.D. 49.5), and AI/FO; 359.1 (S.D. 11.0).

#### MVPA

##### First-level model for concurrent TMS-fMRI task

To obtain estimated activation patterns for MVPA, we estimated a GLM for each participant with SPM5 using the preprocessed images from Session 2. We estimated the activity associated with the two colours and two forms of the objects, using correct trials only. Each trial contributed to the estimation of two beta values: the relevant feature (*green* or *blue* in the colour task, and *cuby* or *smoothy* in the form task) and the irrelevant feature (*cuby* or *smoothy* in the colour task, and *green* or *blue* in the form task), for the Control and Active trials separately (8 regressors per block). To account for trial by trial variation in reaction time (Todd, Nystrom, & Cohen, 2013), trials were modelled as events lasting from stimulus onset until response (Henson, 2007; Grinband, Wager, Lindquist, Ferrera, & Hirsch, 2008; Woolgar, Golland, & Bode, 2014) convolved with the hemodynamic response of SPM5.

##### ROI analysis

We implemented MVPA using the Decoding Toolbox (Hebart, Görgen, & Haynes, 2015) which wraps the LIBSVM library (Chang & Lin, 2011). For each participant and ROI (MD regions, stimulated dlPFC, LOC, V4, early visual cortex), a linear support vector machine was trained to decode colour (green vs. blue) and form (cuby vs. smoothy) when relevant (e.g., cuby vs. smoothy in form task) and irrelevant (e.g., cuby vs. smoothy in colour task) under the two separate TMS conditions (Control or Active) resulting in 8 separate classification schemes. In total, there were 16 blocks for each participant: 8 with colour relevant, and 8 with form relevant. Since TMS trials were intermingled, half of the trials in these 8 blocks contributed to the classification in the Control condition, and half contributed to classification in the Active condition. Each condition (e.g. cuby when relevant under Active TMS) consisted of 27.6 trials (correct only – on average across participants) with a S.D. of 3.2.

For each classification scheme, we used a leave-one-out 8-fold splitter whereby the classifier was trained using the data from 7 out of the 8 blocks and subsequently tested on its accuracy at classifying the unseen data from the remaining block. This procedure was repeated iterating over all possible combinations of training and testing blocks. The accuracies were then averaged over iterations. This was repeated for each classification scheme, participant and ROI separately. We did not use feature selection or dimensionality reduction.

We entered the classification scores for the MD regions into a four factor ANOVA with factors *TMS* (Control, Active), *Feature* (Colour, Form), *Relevancy* (Relevant, Irrelevant) and *Region* (Left dlPFC, Left AI/FO, Right AI/FO, ACC/pre-SMA, Left IPS, Right IPS). For the remaining ROIs (right dlPFC, LOC, V4, early visual cortex), since they do not form a single network, we conducted separate ANOVAs with factors *TMS* (Control, Active), *Feature* (Colour, Form), and *Relevancy* (Relevant, Irrelevant). Significant interactions were followed up with post hoc analyses. ANOVA main effects and interactions are reported with two-tailed p-values. We also report 95% confidence intervals and effect size; *partial eta squared* for ANOVA calculated in SPSS (IBM Corp. Released 2016. IBM SPSS Statistics, Version 24.0), and Cohen’s *d* calculated in JASP v0.9 (Team, 2018). We applied Bayes Factor (BF) analyses using a default uniform prior and Markov chain Monte Carlo settings in JASP v0.9 (Team, 2018) to interpret all null effects: BF > 3 indicates evidence for experimental hypothesis, and BF < 1/3 indicates evidence for the null hypothesis (Dienes, 2011; Wagenmakers et al., 2017)). Where outliers were present (± 3 SD), we performed a log transformation (Log10) on the raw classification output prior to conducting ANOVAs and post hoc analyses. This was the case for classification in the MD network.

Classification accuracies were compared against chance, where appropriate, using a two-step permutation test (Stelzer, Chen, & Turner, 2013). For this, we exhaustively permuted the class labels for each decoding analysis (128 unique combinations) and trained and tested the classifier on each permutation. Next we built a group-level null distribution for each condition by sampling (with replacement) 10,000 times from the set of participants x 128 permutation results. Finally, we calculated the probability (*p*) of the observed decoding accuracy given the null distribution, in which *p* = (*k*+1) / (*n*+1) where *k* is the number of permutations in the group null with equal or higher accuracy to the actual value and *n* is the number of all permutations in the group null.

##### Searchlight analysis

To identify any additional brain regions coding task-related information under Control and Active TMS, we carried out an exploratory classification analysis using a roaming spotlight (Kriegeskorte, Goebel, & Bandettini, 2006). In addition to revealing any large effects that our ROI analysis missed, this analysis could potentially be useful to identify candidate ROIs for future studies. For each participant, we extracted data from a spherical ROI (radius 10mm^1^) centred in turn on each voxel in the brain. A linear support vector machine was trained and tested as before, using data from each sphere, and the classification value for that sphere was assigned to the central voxel yielding whole brain classification maps for each individual.

To combine data across individuals, individual-subject classification accuracy maps were normalised and subsequently smoothed using an 8mm FWHM Gaussian kernel. In addition to a repeated-measures ANOVA including all experimental conditions (factors: *TMS, Feature, Relevancy*), mimicking our ROI analyses, paired t-tests were conducted to directly compare coding between Active and Control TMS under the two relevancy conditions. Classification accuracy was compared to chance at the group level using a one-sample t-test against chance (50%). Since we ran this exploratory analysis to identify additional regions outside of our *a priori* ROIs where information coding was modulated by TMS, we used a lenient voxelwise threshold of *p <* 0.0001 and corrected for multiple comparisons at the cluster-level using FWE (*p* < 0.05). We identified the anatomical locations of significant clusters using the Brodmann and AAL templates of MRICroN (Rorden, 2007) and the Harvard-Cortical and subcortical structural atlases of FSL (Jenkinson, Beckmann, Behrens, Woolrich, & Smith, 2012).

## Results

### Behavioural data

We compared behavioural data for stimuli in which colour and form mapped onto the same button-press response (congruent) to those where the two stimulus dimensions indicated different button-press responses (incongruent), using three-way repeated measures ANOVAs, with factors *TMS* (Control, Active), *Feature* (Colour, Form), and *Congruency* (Congruent, Incongruent). For reaction time (RT) data (correct trials only, Figure 3, left panel), there was a significant interaction between *TMS* and *Congruency* (*F*(1,19) = 4.78, *p* = 0.04, *η*^*2*^_*p*_ = 0.2). Post-hoc paired t-tests showed that participants were significantly faster on congruent (518ms) than incongruent trials (535ms; *t*(19) = 4.78, M_diff_ = 16.5 [95% CI: 9.27, 23.7], p < 0.001, *d* = 0.34) in the Control TMS condition, but in the Active condition there was no significant difference between congruent (532ms) and incongruent (530ms) trials (*t*(19) = 0.36, M_diff_ = 2.34 [95% CI: -11.3, 16.1], *p* = 0.73, *d* = 0.04), with Bayes analysis indicating evidence for the null hypothesis of no effect (BF_10_ = 0.25). Participants were also significantly faster in the Control compared to the Active TMS condition on congruent trials (*t*(19) = 2.68, M_diff_ = 13.9 [95% CI: 2.12, 25.9], *p* = 0.02, *d* = 0.55), but there was no significant difference in RT performance between the Control and Active TMS conditions on incongruent trials (*t*(19) = 0.89, M_diff_ = 4.86 [95% CI: -6.5, 16.2], *p* = 0.38, *d* = 0.21, BF_10_ = 0.33). No other main effects or interactions were significant (all *p*s > 0.11, all BF_10_ < 0.99).

For accuracy data (percent correct; Figure 3, right panel) there was a main effect of *Congruency* (*F*(1,19) = 11.58, M_diff_ =3.23 [95% CI: 1.26, 5.29], *p* = 0.003, *η*^*2*^_*p*_ = 0.38), reflecting more accurate performance on congruent (87.2%) relative to incongruent (84%) trials overall. No other main effects or interactions were significant (all *p*s > 0.23, all BF_10_ < 0.48). Therefore, Active TMS had a significant effect on RT, and no detectable effect on accuracy.

### The effect of TMS on overall activation levels

We conducted a whole-brain mass-univariate analysis to examine whether overall activation levels in different brain regions were affected by Active TMS to right dlPFC (Figure 4; Table 1). Relative to Control TMS, Active TMS increased BOLD response in clusters of voxels in and around the left temporal cortices (Heschl’s gyrus, superior temporal gyrus), likely reflecting the difference in auditory stimulation between the two TMS conditions. We also observed increased BOLD in clusters in/around the frontoparietal MD network (right ACC, left dlPFC) and visual cortices (left extrastriate cortex, right primary visual cortex), under Active TMS, demonstrating long-range effects of dlPFC stimulation. The reverse contrast (Control > Active) showed no significant clusters.

**Table 1:** Peak coordinates for univariate contrast Active > Control TMS. The results were thresholded at *p <* 0.0001 (FWE correction of *p* < 0.05 at cluster level). N=20.

| Contrast | Cluster | Hemisphere | Peak coordinates |  |  | Brodmann area | Cluster size | t | FWE (p) |
| --- | --- | --- | --- | --- | --- | --- | --- | --- | --- |
|  |  |  | x | y | z |  |  |  |  |
| Univariate: Active > Control | dorsolateral prefrontal cortex | left | -40 | 38 | 10 | 45 | 225 | 7.32 | <0.0001 |
|  | primary visual cortex |  |  |  |  |  |  |  |  |
|  | extending into extrastriate |  |  |  |  |  |  |  |  |
|  | cortex | right | 14 | -78 | 0 | 17/18 | 416 | 7.13 | <0.0001 |
|  | Heschl's gyrus | left | -42 | -22 | 12 | 48 | 236 | 6.83 | <0.0001 |
|  | superior temporal gyrus | left | -44 | 2 | 6 | 13 | 42 | 6.41 | =0.046 |
|  | anterior cingulate cortex | right | -8 | 42 | 18 | 32 | 57 | 6.23 | =0.019 |
|  | superior temporal gyrus | left | -38 | -4 | -6 | 22 | 197 | 5.77 | <0.0001 |
|  | superior temporal gyrus | right | 38 | -12 | -8 | 22 | 56 | 5.75 | =0.02 |
|  | extrastriate occipital | left | -14 | -62 | -2 | 18 | 61 | 5.53 | =0.015 |

### The effect of TMS on information coding

#### Unstimulated MD regions

Next we turned to our main question, which was whether TMS to right dlPFC affected *information coding*, and whether this modulation differed for information that was relevant vs irrelevant to the current task. For this, we examined coding of object colour (green vs blue) and form (cuby vs smoothy) when relevant (e.g., cuby vs smoothy in form task) and irrelevant (e.g., cuby vs smoothy in colour task) under the two separate TMS conditions (Control and Active). First, we examined information coding in the unstimulated MD network. Figure 5 shows the classification accuracies for the different MD regions for relevant (upper panel) and irrelevant (lower panel) information separately. We entered these classification accuracies into an ANOVA with factors *TMS, Feature, Relevancy*, and *Region.* There was a significant interaction between *TMS* and *Relevancy* (*F*(1,19) = 5.58, *p* = 0.03, *η*^*2*^_*p*_ = 0.23) indicating that that TMS had a differential impact on coding of relevant and irrelevant information. Post-hoc paired t-tests revealed that relevant information coding was significantly reduced under Active TMS (51.3%) compared to Control TMS (56.8%; *t*(19) = 2.21, M_diff_ =5.42 [95% CI: 0.67, 10.2], *p* = 0.04, *d =* 0.53). However, for irrelevant information coding there was no significant difference between Control (54.2%) and Active (55%) TMS conditions (*t*(19) = 0.28, M_diff_ =0.79 [95% CI: -0.51, 6.64], *p* = 0.78, *d =* -0.06), and Bayes analyses indicated evidence for the null hypothesis of no effect (BF_10_ = 0.08). The ANOVA also revealed a main effect of *Feature* (*F*(1,19) = 6.21, M_diff_ =5.04 [95% CI: 0.85, 9.21], *p* = 0.02, *η*^*2*^_*p*_ = 0.25) driven by overall stronger coding of colour (56.9%) compared to form (51.8%), but this baseline effect did not interact with any other factors. No other main effects (all *p*s > 0.36, all BF_10_ < 0.69) or interactions (all *p*s > 0.13, all BF_10_ < 0.12, task*relevancy interaction *p* = 0.09 and BF_10_ = 6.44) were significant.

The significant effect of TMS on decoding of relevant information is only interpretable if coding in one or more of the TMS conditions is significantly above chance. Therefore, we compared classification accuracies against chance, using a two-step permutation test. We found that relevant information was significantly coded under Control TMS (permutation test, *p* < 0.001) but was not under Active TMS (by permutation, *p* = 0.35, BF_10_ = 0.18). Therefore, the significant effect of TMS on relevant information coding is interpretable: TMS to right dlPFC significantly reduced coding of relevant stimulus information in the rest of the MD system. The interaction further specifies that this effect was larger than any effect on *irrelevant* information coding, which was not detected.

#### Right dlPFC (stimulated site)

The right dlPFC was the target for stimulation so we analysed it separately from the rest of the MD system (Figure 5, right panels). It is typical not to see effects at the stimulation site with our setup (Bestmann, Ruff, Driver, & Blankenburg, 2008), one reason for this being that the TMS coil may shield this part of the cortex from MR excitation and readout. Although the trend appeared to be in the same direction as the rest of the MD system, our ANOVA (factors *TMS, Feature*, and *Relevancy*) detected no significant main effects or interactions (all *p*s > 0.2, all BF_10_ < 0.46).

#### Visual cortices

Next we examined top-down effects in the visual cortex. The trend in the early visual cortex ROI (defined from individual-subject localiser data; Figure 6, left panel) was for a decrease in relevant *and* an increase in irrelevant information coding, but the statistical analysis did not show any significant main effects or interactions (all *p*s > 0.06, all BF_10_ < 0.63).

In LOC (defined from individual-subject localiser data; Figure 6, centre panel), we observed a different pattern. Here, there was a significant main effect of *TMS* (*F*(1,19) = 10.1, M_diff_ =5.15 [95% CI: 1.75, 8.55], *p* = 0.005, *η*^*2*^_*p*_ = 0.34) with stronger coding for Control than for Active TMS, without a *TMS*\**Relevancy* interaction (*F*(1,19) = 0.03, *p* = 0.86, *η*^*2*^_*p*_ = 0.002, BF_10_ = 0.008). These data align with a general top-down effect of dlPFC activity on all stimulus processing, irrespective of relevancy. However, this main effect was modulated by a significant interaction between *TMS* and *Feature* (*F*(1,19) = 6.28, *p* = 0.02, *η*^*2*^_*p*_ = 0.25), which reflected a stronger effect of TMS on colour coding (Control 57.9%, Active 48.1%, post-hoc paired t-test *t*(1,19) = 4.21, M_diff_ = 9.87 [95% CI: 4.97, 14.78], *p* < 0.0001, *d =* 0.94) than on form coding (Control 53.4%, Active 52.9%, *t*(1,19) = 0.17, M_diff_ = 0.44 [95% CI: -5.05, 5.92], *p* = 0.87, *d =* 0.04, BF_10_ = 0.23). This may reflect the overall tendency, seen also in the MD regions, for colour to be coded more strongly than form in our task. No other main effects or interactions were significant (all *p*s > 0.47, BF_10_ < 0.35).

In colour-responsive cortex (V4, defined by coordinates from the literature; Figure 6, right panel) we saw a similar pattern. There was again a significant main effect of *TMS* (*F*(1,19) = 9.72, M_diff_ =5.18 [95% CI: 1.71, 8.67], *p* = 0.006, *η*^*2*^_*p*_ = 0.23) with stronger coding for Control than for Active TMS, and no *TMS*\**Relevancy* interaction (*F*(1,19) = 0.01, *p* = 0.91, *η*^*2*^_*p*_ = 0.001, BF_10_ = 0.22). There was also, again, a significant *TMS*Feature* interaction (*F*(1,19) = 5.61, *p* = 0.03, *η*^*2*^_*p*_ = 0.23), reflecting a stronger effect of TMS on colour than on form. Post-hoc paired ests showed a significant effect of TMS on colour coding (Control 59.6%, Active 50.2%, post-hoc paired t-test *t*(1,19) = 4.17, M_diff_ = 9.37 [95% CI: 4.67, 14.1], *p* = 0.001, *d =* 0.93) and no effect on form coding (Control 51.9%, Active 50.9%, *t*(1,19) = 0.38, M_diff_ = 0.99 [95% CI: -4.44, 6.43], *p* = 0.71, *d =* 0.09, BF_10_ = 0.25). No other main effects or interactions reached significance (all *p*s > 0.06, all BF_10_ < 1.00).

Overall, right dlPFC-TMS had a significant effect on information coding in higher visual cortex ROIs (LOC, V4), but there was no evidence that this effect differed for relevant and irrelevant information processing.

#### Searchlight analysis

We conducted an exploratory analysis to check for additional regions in which coding was affected by TMS to the right dlPFC by performing decoding analyses across the whole-brain using a roaming searchlight (Kriegeskorte et al., 2006). The advantage of this approach is that is it free from *a priori* spatial hypotheses, meaning we can potentially identify additional regions missed by the ROI approach, but it has a lack of power relative to ROI analyses, given the large number of comparisons it entails.

We performed searchlights comparing coding to chance in each of the relevancy and TMS conditions separately, and then compared these maps to one another. Under Control TMS, significant coding of relevant information was seen across the brain including in/around the MD network and visual cortices (Figure 7 upper panel, Table 2). For irrelevant coding, five large clusters survived correction at the cortical and subcortical level under Control TMS (Figure 7 lower panel, Table 2). For both conditions, no clusters survived correction under Active TMS.

**Table 2:** Peak coordinates for whole-brain searchlights. Relevant information under Control (upper panel), and irrelevant information under Control TMS (lower panel). The results were thresholded at *p <* 0.0001 (FWE correction of *p* < 0.05 at cluster level). N=20.

| Contrast | Cluster | Hemisphere | Peak coordinates |  |  | Brodmann area | Cluster size | t | FWE (p) |
| --- | --- | --- | --- | --- | --- | --- | --- | --- | --- |
|  |  |  | x | y | z | at peak |  |  |  |
| Relevant (Control condition) | peak in precuneus/cingulate gyrus. cluster extends to cerebellum/lingual gyrus/intraparietal sulcus/precentral gyrus/anterior cingulate/anterior insula/middle and inferior frontal gyrus | peak in left but cluster extends bilaterally |  |  |  |  |  |  |  |
|  |  |  | -12 | -56 | 8 | 17 | 39083 | 12.77 | <0.0001 |
|  | precentral gyrus | right | 54 | 6 | 26 | 6 | 468 | 6.45 | =0.002 |
|  | supramarginal gyrus | left | -40 | -34 | 28 | 48 | 872 | 6.15 | <0.0001 |
|  | middle frontal gyrus | right | 34 | 28 | 20 | 48 | 138 | 5.08 | =0.038 |
| Irrelevant (Control condition) | peak in precentral gyrus. cluster extends to precuneus, anterior cingulate, lateral occipital complex, intraparietal sulcus, superior parietal lobule and middle frontal gyrus | peak in right but cluster extends bilaterally |  |  |  |  |  |  |  |
|  |  |  | 0 | -32 | 60 | nearest is 4 | 7815 | 8.99 | <0.0001 |
|  | peak in amygdala. cluster extends to precuneus, hippocampus, insula, putamen and lateral occipital complex | peak in left but cluster extends bilaterally | -20 | -4 | -24 | 28 | 12456 | 7.16 | <0.0001 |
|  | frontal orbital cortex | right | 30 | 32 | -8 | 47 | 172 | 6.18 | =0.03 |
|  | paracingulate gyrus | right | 8 | 46 | 6 | 32 | 449 | 5.73 | =0.004 |
|  | paracingulate gyrus | right | 10 | 32 | 30 | 32 | 269 | 5.68 | =0.013 |

The direct comparison of these maps revealed stronger coding of relevant information under Control compared to Active TMS in seven clusters, including in and around the ACC, left dlPFC, as well as in occipital, frontal and temporal cortices (Table 3, Figure 7; upper panel). There were no significant clusters for the reverse contrast (Active > Control). For irrelevant information (Control > Active), there were several significant clusters including in occipital and temporal cortices (Table 3, Figure 7; lower panel). There were no significant clusters for the reverse contrast (Active > Control).

**Table 3:** Peak coordinates for whole-brain searchlights. Relevant information under Control > Active (upper panel), and irrelevant information under Control > Active (lower panel). The results were thresholded at *p <* 0.0001 (FWE correction of *p* < 0.05 at cluster level). All significant clusters had regions that were also significantly coded against chance (depicted in Figure 7). N=20.

| Contrast | Cluster | Hemisphere | Peak coordinates |  |  | Brodmann area<br>at peak | Cluster size | t | FWE (p) |
| --- | --- | --- | --- | --- | --- | --- | --- | --- | --- |
|  |  |  | x | y | z |  |  |  |  |
| **Relevant information**<br>(Control > Active) |  | peak in right but cluster extends |  |  |  |  |  |  |  |
|  | superior frontal gyrus extending towards anterior cingulate | bilaterally | 12 | 10 | 64 | 6 | 817 | 8.82 | <0.0001 |
|  | middle temporal gyrus | left | -62 | -28 | -18 | 20 | 298 | 7.81 | =0.004 |
|  | lateral occipital complex extending to precuneus | right | 34 | -56 | 40 | 40 | 762 | 6.88 | <0.0001 |
|  |  | peak in left but cluster extends |  |  |  |  |  |  |  |
|  | anterior cingulate gyrus | bilaterally | -4 | 20 | 28 | 24 | 835 | 6.77 | <0.0001 |
|  | cerebellum | left | -8 | -50 | -18 | 19 | 231 | 6.39 | =0.009 |
|  | peak in temporal occipital fusiform. clusters extends to cerebellum and parahippocampal gyrus | right | 34 | -38 | -18 | 37 | 749 | 6.12 | <0.0001 |
|  | peak in lingual gyrus. cluster extends to occipital fusiform and intracalcarine cortex. | bilateral | 0 | -78 | -14 | 14 | 308 | 5.52 | =0.004 |
| **Irrelevant information**<br>(Control > Active) | peak in occipital fusiform gyrus. cluster extends to occipital pole and lateral occipital complex | left | -22 | -84 | -4 | 18 | 811 | 7.09 | <0.0001 |
|  | peak in temporal fusiform extending into parahippocampal gyrus | left | -40 | -16 | -20 | 20 | 531 | 6.95 | =0.001 |
|  | thalamus | right | 12 | -8 | -4 | nearest is 48 | 202 | 6.92 | =0.016 |
|  | occipital fusiform gyrus | left | -40 | -58 | -16 | 37 | 385 | 6.51 | =0.003 |
|  | cingulate gyrus | left | -4 | -6 | 42 | 24 | 204 | 6.14 | =0.016 |
|  | heschl's gyrus | right | 36 | -20 | 16 | 48 | 325 | 5.84 | =0.005 |
|  | lingual gyrus | bilateral | 6 | -74 | -2 | 17 | 208 | 5.77 | =0.015 |
|  | supramarginal gyrus | left | -46 | -28 | 22 | 48 | 125 | 5.7 | =0.039 |
|  | precuneus | right | 10 | -36 | 62 | nearest is 4 | 126 | 5.09 | =0.039 |

Finally, for comparison of all our conditions, we analysed the data with a repeated measures ANOVA (factors: *TMS, Feature, Relevancy*). This showed no significant clusters for either the three-way, or the two-way interactions. Significant clusters were observed in the cingulate gyrus, precuneus, visual cortices, cerebellum, subcallosal cortex, superior temporal gyrus, brain stem and post-/pre-central gyrus for the main effect of *TMS* (Control > Active; Figure 8, upper panel), indicative of a drop of information under Active TMS, and *Feature* (Colour > Form; Figure 8, lower panel), reflecting stronger coding of colour than shape, as in the ROI analyses. There were no significant clusters for the main effect of *Relevancy*.

The results of these exploratory analyses indicate that right dlPFC activity supports coding of task-related information across a range of brain regions. However, in concordance with the pre-defined ROI analyses, there was no evidence for a release from suppression following stimulation to right dlPFC. Qualitatively the effect of TMS on relevant information appears to be larger in the resultant maps, however, this interaction did not reach significance as it did in the pre-defined MD network.

## Discussion

We used multivariate analyses of concurrent TMS-fMRI data to causally examine the role of right dlPFC in supporting attentional processing in the brain. In a selective attention task, perturbing right dlPFC affected information coding in the MD network, visual cortices, and other brain areas, commensurate with a role for the right dlPFC in supporting brain-wide information processing. Critically, the data suggest that the role of the dlPFC is primarily in supporting the coding of *task-relevant* information processing. Active TMS impaired MD coding of relevant visual object information (object form or colour when needed for the task), but had no detectable effect on MD coding of the identical visual object information when it was not needed for the task (e.g. an object’s colour when participants were reporting the object’s form). Parametric statistics confirmed that the effect of TMS on relevant information coding was significantly larger than any effect on irrelevant information, and Bayesian analyses suggested evidence for the *absence* of an effect on irrelevant information. These data contribute to our understanding of causal mechanisms giving rise to the prioritisation of task-relevant information in the human brain.

The dlPFC is thought to be crucial for goal-directive behaviour. It is part of a circuit of frontal and parietal brain regions, referred to as the MD regions (Duncan, 2010), task-positive network (Fox et al., 2005), or frontoparietal control system (Vincent, Kahn, Snyder, Raichle, & Buckner, 2008). These areas are known to be engaged by a wide variety of task demands (e.g. Duncan & Owen, 2000; Fedorenko et al., 2013) and have been posited to play a fundamental role in executive function and cognitive control (Corbetta & Shulman, 2002; Cole & Schneider, 2007). Non-human primate research has shown that these regions preferentially code task-relevant information (Rao, Rainer, & Miller, 1997; Freedman et al., 2001; Cromer et al., 2010; Roy et al., 2010; Kadohisa et al., 2013), a result mirrored in human neuroimaging data (Woolgar et al., 2015; Jackson et al., 2017; Jackson & Woolgar, 2018). This selective representation may support preferential coding in other brain regions (Desimone & Duncan, 1995; Miller & Cohen, 2001), as indicated by previous work (Baldauf & Desimone, 2014; Goddard, Carlson, & Woolgar, 2019), yielding a plausible mechanism for a brain-wide focus on relevant information and a key component of cognitive control. For example, Baldauf and Desimone (2014) combined magnetoencephalography and fMRI to show that the inferior frontal junction appears to direct object-based attentional inputs to the inferior-temporal cortex. However, these previous data are not sufficient to establish a causal role. In parallel, findings from offline TMS (Olk, Peschke, & Hilgetag, 2015) and patients with chronic focal lesions (e.g. Woolgar, Bor, & Duncan, 2013; Woolgar, Duncan, Manes, & Fedorenko, 2018), suggest a causal role for the MD network in attention and higher cognition, but do not establish the neural mechanisms by which it is achieved. The current work bridges this gap by demonstrating a causal role for dlPFC in supporting the representation of task-relevant information in frontoparietal cortex and other, anatomically distributed, brain regions.

The data further suggest that the mechanism by which dlPFC supported selective processing in our task was primarily through supporting processing of task-relevant information, rather than through suppressing irrelevant information, contributing to a key debate in this field. The dlPFC has previously been suggested to be involved in inhibitory mechanisms (e.g. Knight et al., 1999; Shimamura, 2000; Aron, 2007; Hampshire & Sharp, 2015), leading to the prediction that disrupting dlPFC with TMS should cause a “release from inhibition” for distracting (suppressed) information. This would have been seen as an increase in coding of the irrelevant information. Such a result would also be predicted by the biased-competition framework (Desimone & Duncan, 1995; Hampshire & Sharp, 2015), particularly for visual cortex, based on reduced local competition from the downregulated relevant information. Our data, however, fail to find evidence for either of these accounts, with our Bayes analyses indicating evidence for no effect of TMS on irrelevant information coding (MD regions), and a *reduction* in coding across relevancy conditions in higher visual cortex (LOC and V4 ROIs) and searchlight clusters in/around cingulate gyrus, precuneus, visual cortices, cerebellum, subcallosal cortex, superior temporal gyrus, brain stem and post-/pre-central gyrus. No regions showed an increase in irrelevant information by TMS in the ROI or searchlight analyses. The extent to which we can generalise a finding from one type of task to another is, of course, unknown. It may be possible to see release from suppression with more data, or with a technique with higher temporal resolution, or in a more difficult task that places the different sources of information in more direct competition. Nonetheless the interaction between the effect of TMS and relevancy in the unstimulated MD network specifies that, in this task, the role of right dlPFC was primarily facilitating task-relevant, rather than inhibiting irrelevant, information processing. These findings support major accounts of higher cognitive functions (e.g. Miller & Cohen, 2001; Duncan, 2010) that posit dlPFC as mediating control by facilitating processing of attended information.

TMS also had a significant effect on behaviour, reducing the congruency effect on RT, apparently by reducing the advantage for congruent trials. It is challenging to map this result directly to the neural data, because whilst the neural data separates coding of relevant and irrelevant information, the behavioural data in both congruent and incongruent conditions (presumably) reflect a combination of both. One possibility is that TMS reduced the person’s ability to benefit from congruency because it interfered with processes involved in recognising or responding to congruency signals, but this is not something we could decode with this design. Nonetheless, part of the promise of using concurrent TMS-fMRI with cognitive tasks is the possibility of relating neural activity to behaviour, and future work may capitalise on this by using simpler designs where both neural data and behaviour can be analysed for the same information, or, for example, by interrogating the information content on behavioural errors (Woolgar, Dermody, Afshar, Williams, & Rich, 2019).

The present findings showed a disruptive effect of stimulation, with decreased coding of task information under the Active condition coinciding with a reduced advantage on congruent trials as evident from the RT data. The relationship of these data to the univariate contrast showing that overall BOLD was increased under the Active condition is unclear. Whilst an increase in BOLD could be interpreted as a facilitatory response to stimulation, numerous factors could have led to differences between the Active and Control stimulation conditions here. For example, there was increased activation in several auditory regions under Active TMS, presumably reflecting differences in acoustic noise between the Active and Control conditions. A further note is that overall BOLD level is derived from a combination of inhibitory and excitatory signals, thus it is problematic to interpret an increase under Active TMS as a purely facilitative response. Distinctions between multivariate and univariate analyses are not uncommon (Davis et al., 2014) and we argue that in this case multivariate analyses may be more informative, as they provides us with evidence relating to the integrity of task-related information processing. Future work is still needed to understand the relationship between overall BOLD and information decoding.

One limitation of our setup was that the placement of the TMS coil may have shielded the targeted cortex from MR excitation and readout, giving relatively poor signal under the coil. This meant that we could not independently confirm that we stimulated our precise target of interest. Moreover, since neuronavigation was performed outside of the scanner room and the coil orientation and position occasionally had to be adjusted slightly to accommodate the MR coil, TMS targeting was likely imperfect. However, given that the magnetic field of TMS is somewhat diffuse (Deng, Lisanby, & Peterchev, 2013), and that the area of dlPFC activated as part of the MD network is actually relatively large (e.g. from Duncan and Owens’ (2000) template definition of right dlPFC, the average ROI size for analysis here was 685.6 voxels across participants), we suggest that these small adjustments are unlikely to have a noticeable effect. For future work, we are encouraged by the recent developments of in-MR and in-bore neuronavigation technology (Localite) and dedicated TMS-fMRI high sensitivity coil arrays (Navarro de Lara et al., 2015) which give excellent resolution directly under the coil (∼5% BOLD signal change (de Lara et al., 2017) compared to previous setups with ∼0.5% signal change directly underneath the coil (e.g. Bestmann, Baudewig, Siebner, Rothwell, & Frahm, 2003)), increasing the accuracy and confidence in target ROI stimulation.

In the present study, we observed modulated coding of task-related information across the frontoparietal network, as well as long-range effects to other brain networks. The effect of TMS on the coding of relevant information appeared to be more widespread than the modulation of irrelevant information, but the searchlight analysis did not reveal the same TMS*Relevancy interaction that was observed in the pre-defined MD network. This may be due to a relative lack of power in the searchlight analysis or may reflect a more general top-down effect of dlPFC-TMS which affects representation of both types of task information. Areas where coding of relevant information was reduced under Active TMS (and relevant information was coded above chance under the Control condition) included occipital cortices (peak in lingual gyrus), cerebellum, LOC (extending to precuneus), and the MD regions (ACC/pre-SMA and left dlPFC). Comparatively, regions where coding of irrelevant information was reduced under Active TMS (and irrelevant information was coded above chance under the Control condition) included visual areas (occipital fusiform, LOC), cerebellum, temporal cortices, precuneus and thalamus. The data emphasise that perturbation of right dlPFC causes top-down modulation of information in distant regions, underscoring the long-range connectivity associated with frontoparietal cortex.

With these data, we address the longstanding debate of whether the dlPFC exerts top-down control via the enhancement of task-relevant information, and/or via the inhibition of task-irrelevant information. The findings provide causal support for only the former mechanism, with perturbation of right dlPFC with TMS specifically disrupting coding of *relevant*, as opposed to the equivalent *irrelevant*, information in the unstimulated frontoparietal network. Exploratory analyses suggested that disruption to dlPFC also affected coding of both relevant and irrelevant information in discrete distal regions, but there was no evidence for a release from suppression of irrelevant information anywhere in the brain. The data provide strong causal evidence in support of major theories implicating prefrontal cortex in executive control and suggest that the primary mechanism for control by right dlPFC is in biasing processing *towards* information that is relevant to our behaviour.

## Acknowledgements

This work was funded by an ARC Centre of Excellence in Cognition and its Disorders Student Exchange Grant and a Macquarie University Department of Cognitive Science Postgraduate Grant awarded to JJ. Scanning acquisition was funded by the Centre for Integrative Neuroscience and Neurodynamics, University of Reading. JJ was supported by an International Macquarie University Research Excellence Scholarship from Macquarie University, an EMCG grant (MR/N026969/1) and MRC (U.K) intramural funding SUAG/052/G101400. ANR and AW are supported by ARC Discovery Project grants (DP170101840, and previously DP12102835). AW was a recipient of an ARC Future Fellowship (FT170100105) and is now supported by MRC (U.K) intramural funding SUAG/052/G101400. We thank Hans Op de Beeck for providing the scripts used to generate the stimuli.

## Conflicts of Interest Statement

The authors declare no competing financial interests.

## Footnotes

1 The whole-brain searchlight analysis was originally conducted at 5mm, then changed to 10mm at reviewer request. The effect on relevant coding at 10mm was slightly more extensive but in both cases, there was no indication of a release from suppression.

## Notes

### Competing Interest Statement

The authors have declared no competing interest.

